# Speech and music organize neural activity differently over time in human superior temporal cortex

**DOI:** 10.64898/2026.11.19.767838

**Authors:** Donghui Song, Henyi Rao

**Affiliations:** Department of Psychology, University of Science and Technology of China, Hefei, China

**Keywords:** intracranial EEG, superior temporal gyrus, speech, music, sample entropy, naturalistic auditory processing, fMRI

## Abstract

Speech and music engage overlapping auditory cortex. We asked whether they shape the timing of local activity in ways that regional response strength alone leaves unresolved. In intracranial recordings from 30 participants watching a film with alternating music and speech, 1–10-Hz activity in the superior temporal gyrus (STG) had higher sample entropy during music, indicating that similar short patterns were less likely to persist (mean paired difference = 0.02256, FDR q = 0.00753). The contrast persisted in comparisons of adjacent film blocks and was larger relative to surrogate signals matched for amplitude distribution and approximate power spectrum. Speech elicited higher STG high-frequency power in the intracranial subset and stronger bilateral STG BOLD responses in all 30 participants in the separately acquired fMRI recordings. Together, these findings show that the soundtrack category associated with stronger regional responses differs from the one associated with greater local temporal irregularity. The temporal organisation of local activity therefore adds a dimension to how shared auditory cortex responds to natural sounds.

## Introduction

Speech and music have distinct temporal structures, yet much of the human auditory cortex responds to both. The superior temporal gyrus (STG) represents spectrotemporal and phonetic features of speech and tracks information at syllabic and phonemic timescales (Giroud et al., 2024; Hullett et al., 2016; Mesgarani et al., 2014). Studies of natural sounds show that speech and music recruit overlapping networks while eliciting distinct response preferences within auditory cortex (Norman-Haignere et al., 2015, 2022a; te Rietmolen et al., 2024). Auditory cortical regions also differ in the time over which they integrate sound (Norman-Haignere et al., 2022b, 2025). These findings establish where speech and music engage auditory cortex and how its responses follow sound. They leave open how the local activity itself unfolds when the same region is engaged by each.

Naturalistic film recordings illustrate this gap. The original report on ds003688 described speech-preferred high-frequency activity and condition-sensitive fMRI responses (Berezutskaya et al., 2022a). Subsequent work found that slow intracranial activity tracks the envelopes of both speech and music, while high-frequency tracking is stronger and more anatomically concentrated for speech (Osorio and Assaneo, 2025). Evidence from other recordings also shows that tracking depends on the temporal features of the sound (Kubanek et al., 2013; Zuk et al., 2021). Response magnitude describes how strongly a region responds, and tracking describes how closely its activity follows a stimulus feature. A complementary question is whether the neural time course revisits similar short patterns more often during one kind of listening than the other.

Sample entropy provides a way to characterise this temporal property across recording modalities. In resting-state fMRI, it describes irregularity in relatively slow BOLD fluctuations (Richman and Moorman, 2000; Wang et al., 2014). Lower resting BOLD entropy has been associated with stronger activation and deactivation in task-responsive regions (Lin et al., 2022). In the STG, studies of hearing loss have also linked auditory history to changes in spontaneous activity, including resting BOLD entropy (Chen et al., 2018; Zhu et al., 2026). Intracranial EEG brings the same question of pattern recurrence to the faster timescale of local electrical activity, where it can be studied as speech and music unfold.

Here we ask whether local activity in the human STG revisits short patterns to the same extent during music and speech, and how that temporal organisation relates to the strength of the regional response. If the same cortical territory shows different preferences for response magnitude and pattern recurrence, mapping where and how strongly it responds captures only part of its functional organisation. Understanding both properties can clarify how shared auditory cortex supports distinct forms of natural listening.

## Results

**Fig. 1.**
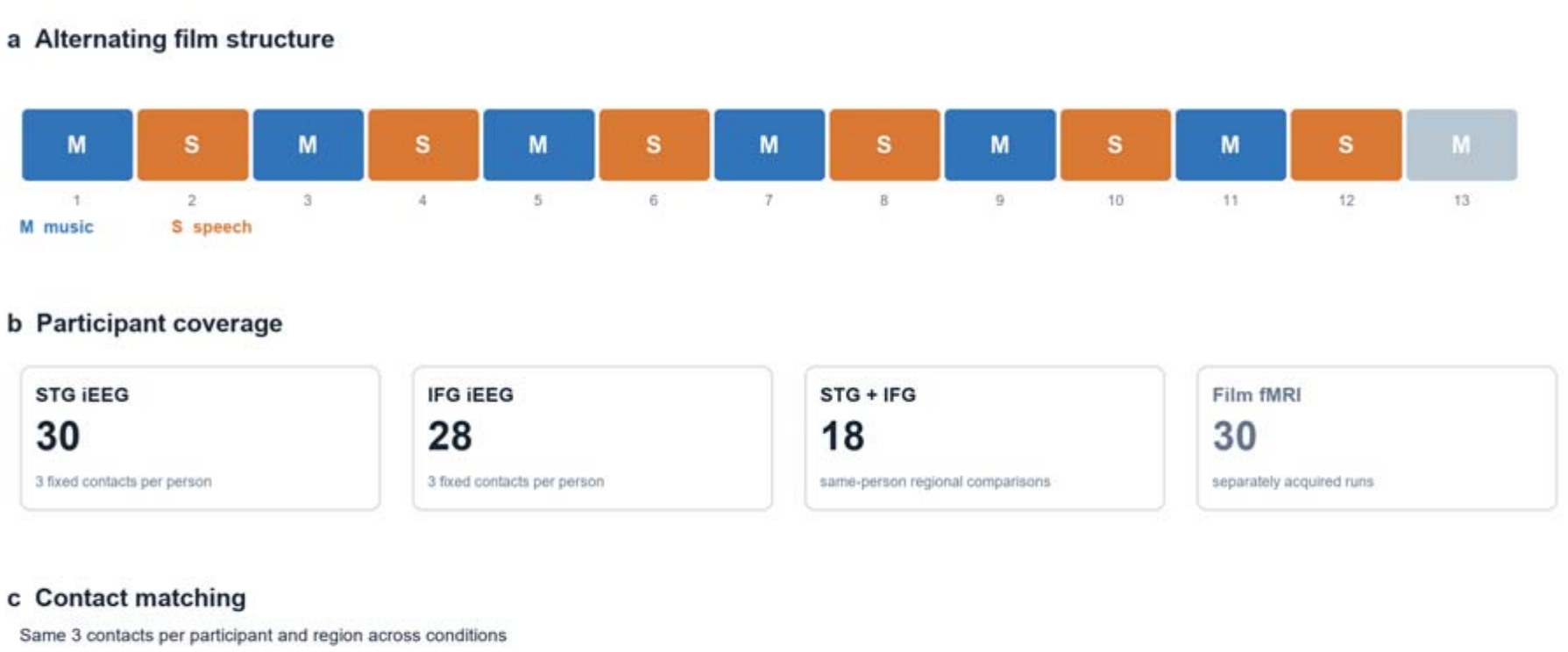
The film alternates music and speech epochs. The primary analysis includes the first six 30-s windows of each condition; the seventh music window is shown in grey and excluded from the balanced primary contrast. Three fixed, anatomically selected left-hemisphere contacts per participant were used for STG (n = 30) or IFG (n = 28). The subset with both regions (n = 18) supports regional interaction analyses and contains 54 STG and 54 IFG contacts. Film fMRI was acquired separately in 30 participants. Blue denotes music and orange denotes speech in the film schematic.

### Low-frequency STG sample entropy is higher during music

Among 30 participants with STG recordings, 1–10-Hz sample entropy was higher during music than during speech (mean paired difference = 0.02256, Cohen’s d_z = 0.571, paired t-test p = 0.00400, FDR q = 0.00753 across three STG contrasts). STG sample entropy was also higher during music than during rest (mean paired difference = 0.04800, paired t-test p = 0.00502, FDR q = 0.00753). The speech–rest contrast and the corresponding IFG contrasts (n = 28) were inconclusive. Including the seventh music block yielded a music–speech difference of 0.02287 (Supplementary Table S1 and Supplementary Fig. S3).

### The STG music–speech contrast persists across surrogate and timing analyses

Relative to IAAFT surrogates, the mean STG music-minus-speech difference in excess sample entropy was 0.03474 (Cohen’s d_z = 1.046, paired t-test p = 3.37 × 10□□, FDR q = 1.01 × 10□□ across three STG contrasts). Including the seventh music block yielded a difference of 0.03527. Surrogate sample entropy alone was higher for speech, and the corresponding IFG excess contrast was inconclusive (Supplementary Table S1).

Comparing speech blocks with the adjacent music blocks yielded a mean STG excess sample entropy contrast of 0.03506 across 30 participants (24 positive, Wilcoxon p = 9.22 × 10□□). With linear and quadratic film time in the model, the estimate was 0.03478 (participant-clustered p = 6.08 × 10□□) (Supplementary Tables S2 and S3 and Supplementary Fig. S3).

**Fig. 2.**
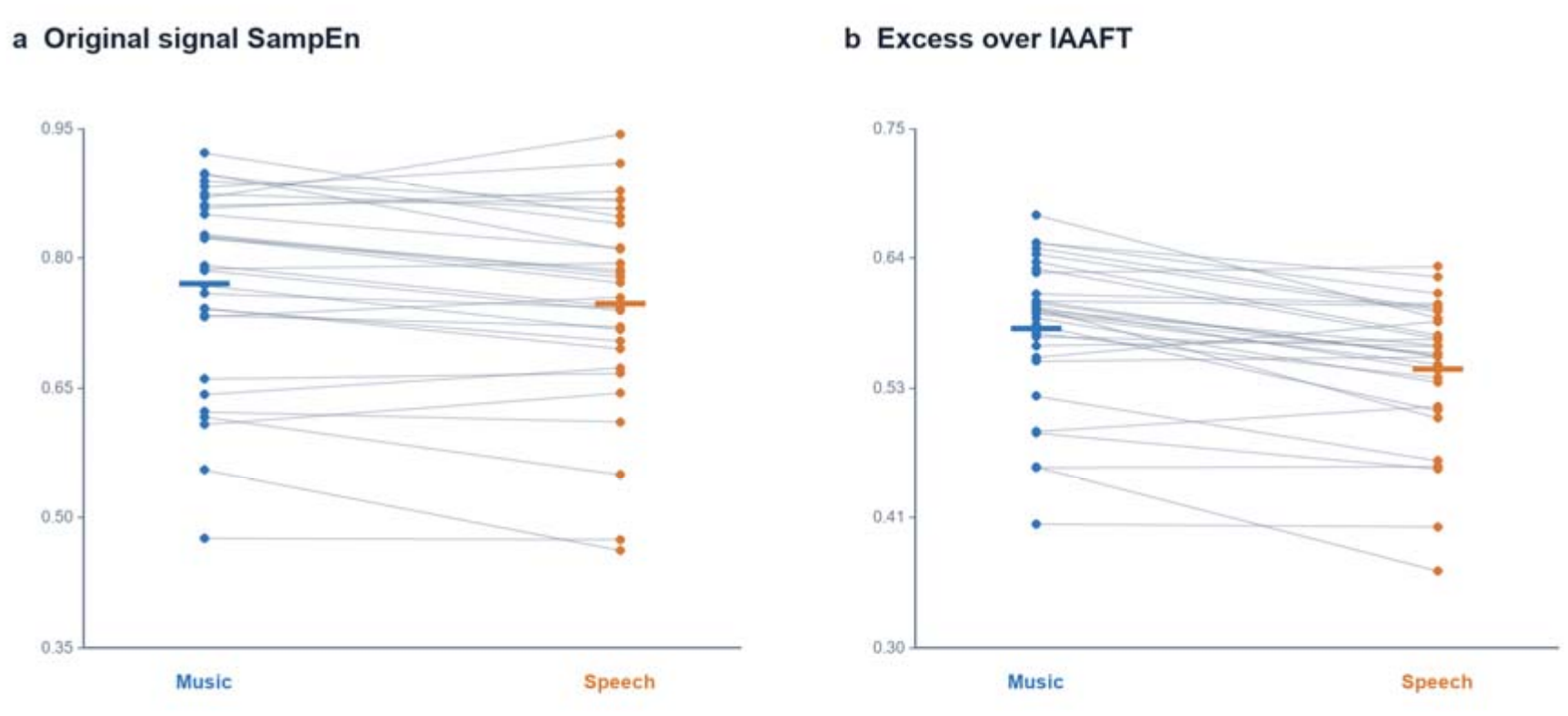
Paired participant values show the STG music–speech contrast for primary 1–10-Hz signal SampEn and excess SampEn relative to IAAFT surrogates. Each line joins condition means from the same participant; horizontal bars mark group means. The mean paired differences are 0.02256 for original SampEn and 0.03474 for excess SampEn; the respective within-STG three-contrast q values are 0.00753 and 1.01 × 10□□. Surrogate entropy alone favoured speech, accounting for the larger excess contrast.

### Frequency-specific sample entropy and power in STG and IFG

Frequency-specific comparisons included 18 participants with recordings in both STG and IFG (Supplementary Figs. S2 and S4).

The STG–IFG interaction in music-minus-speech envelope sample entropy was 0.01681 for alpha (14 of 18 positive, exact sign-flip p = 0.00117, q = 0.00352) and 0.01557 for theta (16 of 18 positive, p = 0.000122, q = 0.000732). The corresponding excess interactions after subtracting surrogate sample entropy were −0.00232 and −0.00241, respectively (Supplementary Table S4).

The high-frequency band (60–100 Hz) envelope sample entropy interaction was 0.02561 (13 of 18 positive, exact sign-flip p = 0.02536, six-band q = 0.05072). The excess interaction was 0.00530 (13 of 18 positive, p = 0.0730).

The STG–IFG interaction in alpha log power was 0.185 in the duration-aligned subset (13 of 17 positive, exact sign-flip p = 0.00475, Supplementary Table S18). In the 18-participant sample, the STG alpha peak above the aperiodic background was 0.516 dB higher during music (14 of 18 positive), with a near-zero difference in IFG. Shifting the alpha boundaries by 0.5 Hz preserved the direction in STG. STG log high-frequency power was higher during speech (music minus speech = −0.166, 14 of 18 in that direction).

**Fig. 3.**
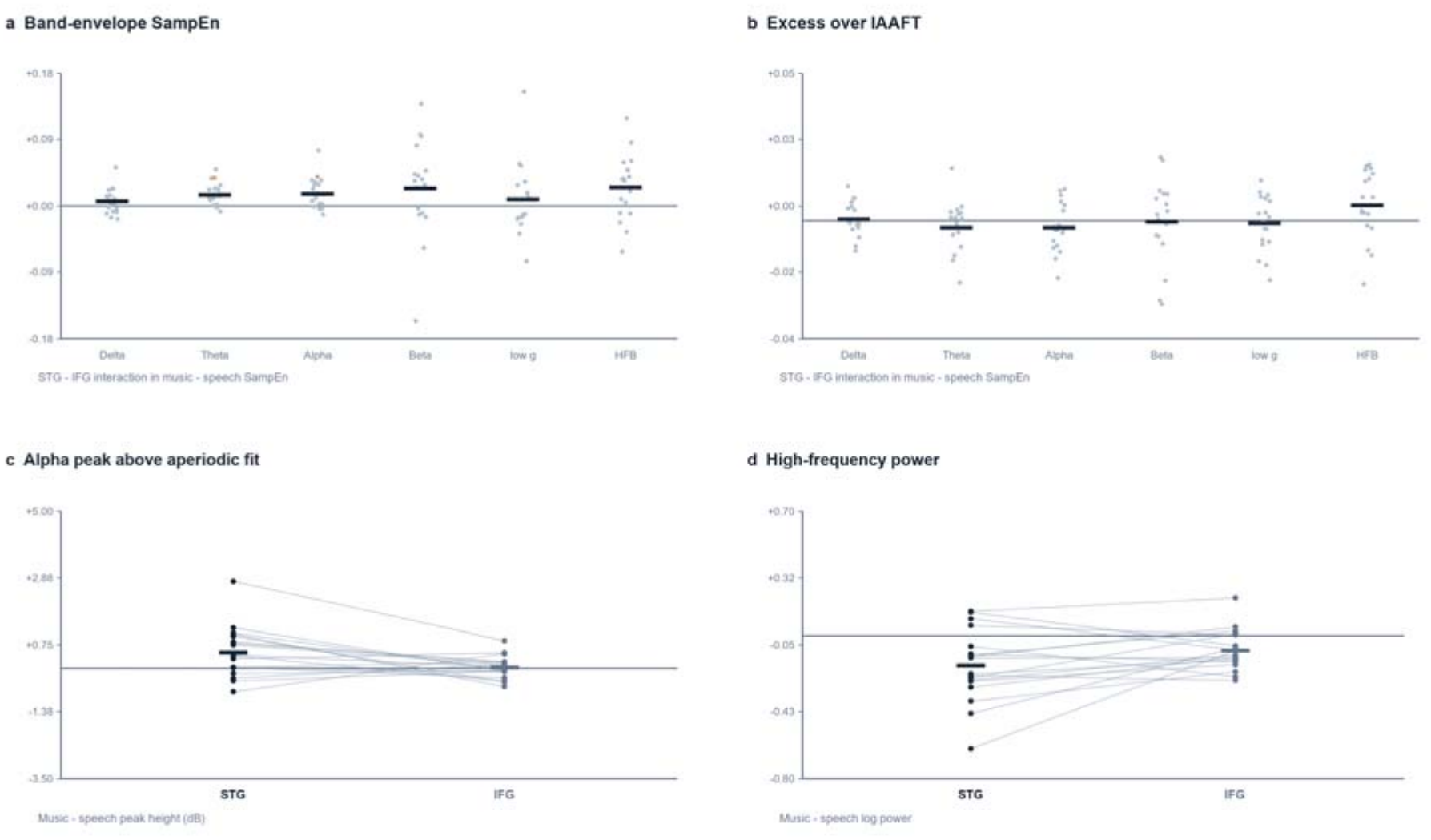
In the 18-person matched STG–IFG subset, panels a and b show participant-level [(STG music − speech) − (IFG music − speech)] interactions for six band-envelope SampEn measures, first in the original signal and then as excess over IAAFT. Dots denote participants, bars the mean, and asterisks six-band false-discovery-rate q < 0.05 within each metric. Panels c and d show STG and IFG music-minus-speech contrasts for alpha peak height above the fitted aperiodic background and log high-frequency power, with within-person lines. Alpha-peak and high-frequency-power units have separate scales. The duration-aligned alpha-power result (n = 17) is reported in the text and Supplementary Table S18.

### STG BOLD activation is higher during speech

Bilateral STG BOLD activation was higher during speech in all 30 participants (mean music-minus-speech difference = −0.751 percentage points). The estimate was similar after adjusting for mean framewise displacement and in the 27-participant lower-motion subset (−0.747 percentage points) (Supplementary Tables S5 and S13).

BOLD sample entropy showed little difference between conditions (music minus speech = −0.00181 for the blockwise mean, Wilcoxon p = 0.952, and −0.00502 for the segmented pooled estimate, p = 0.871) (Supplementary Tables S6 and S14).

Exploratory cross-session correlations are reported in Supplementary Tables S16 and S17.

**Fig. 4.**
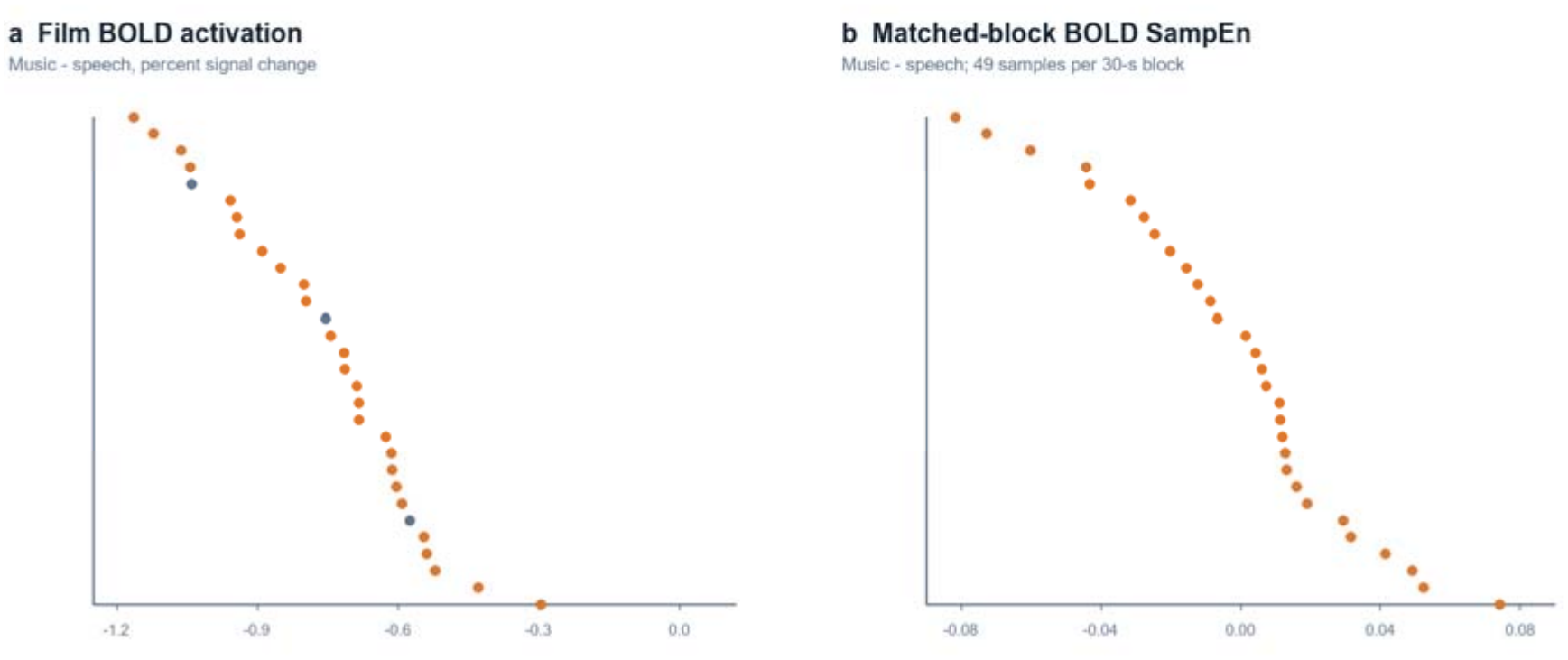
Participant-level STG music-minus-speech BOLD activation estimates for 30 fMRI runs are shown beside matched-block, voxelwise-first BOLD SampEn differences. Grey points in the activation panel identify the three descriptive motion flags. The mean bilateral STG BOLD activation contrast was −0.751 percentage points, indicating greater speech activation in all 30 participants; the 27-participant lower-motion sensitivity estimate was −0.747. The blockwise BOLD SampEn mean difference was −0.00181 (Wilcoxon p = 0.952). The panels use separate horizontal scales; BOLD activation and iEEG were acquired in different sessions.

### Speech annotations are associated with IFG high-frequency power

Among 18 participants, speech blocks with more word onsets had higher IFG high-frequency power (mean within-participant Spearman ρ = 0.505, shared-block permutation p = 0.00833). Exploratory associations with syllable and clause counts were in the same direction (ρ = 0.451 and 0.455, respectively, p = 0.0389 and 0.0333). Associations between the speech annotations and STG sample entropy were inconsistent (Supplementary Fig. S1 and Supplementary Tables S7 and S7b).

## Discussion

Speech and music engage overlapping auditory cortex, yet they favor different properties of the response in the STG. During the film, speech elicited greater high-frequency power and stronger BOLD responses, whereas music showed higher sample entropy in local 1–10-Hz intracranial activity. When short patterns of activity were similar, they were less likely to remain similar as they unfolded during music. The sound that elicits the larger regional response can therefore also elicit a more recurrent local time course.

Previous studies have identified overlapping cortical networks for speech and music alongside populations with distinct response preferences (Norman-Haignere et al., 2015, 2022a; te Rietmolen et al., 2024). The original film study described speech-preferred high-frequency responses (Berezutskaya et al., 2022a). A subsequent analysis showed that slow intracranial activity tracked both soundtrack envelopes, while high-frequency tracking was stronger and more anatomically concentrated for speech (Osorio and Assaneo, 2025). Tracking describes correspondence with an external sound feature. Sample entropy describes whether similar short patterns in the neural signal tend to evolve in similar ways. At a broader spatial scale, Runfola et al. (2025) reported greater interregional manifold-flow complexity during speech. These observations suggest that local pattern recurrence and changes in activity across regions capture different aspects of natural listening.

One explanation for lower STG sample entropy during speech is that successive acoustic landmarks evoke similar local responses. Oganian and Chang (2019) showed that a region of middle STG responds to rapid rises in the speech envelope that provide cues to syllable timing. Giroud et al. (2024) further showed that human auditory cortex tracks syllabic structure in the theta range. If successive landmarks evoke low-frequency waveforms with similar shapes and timing, short patterns could recur even when those landmarks are unevenly spaced. Music also contains salient onsets. The acoustic account therefore predicts that recurrence should vary with the consistency of landmark-locked responses in both categories. One test would match speech and music segments for onset rate and modulation spectrum, measure the similarity of landmark-locked responses, and estimate how much the entropy contrast changes after accounting for those responses.

A second possibility is that linguistic predictions make responses to recurring speech features more consistent. Orepić et al. (2026) found that phonetic population activity in anterior STG aligned with word-level semantic predictions. At the scale of our recordings, a predictive context might constrain the range of local responses to an upcoming word. This hypothesis predicts that the same spoken target will evoke more consistent low-frequency responses after a predictive context than after a neutral context. An acoustic account predicts that varying landmark timing or strength will change recurrence even when linguistic predictability stays constant. Manipulating both factors would test whether context changes the response to a given acoustic landmark.

The spectral results narrow the range of explanations for the local entropy difference. Music showed a more prominent STG alpha peak, while speech elicited greater high-frequency power. The regional interactions in alpha- and theta-envelope sample entropy were attenuated after accounting for IAAFT surrogate entropy. The primary 1–10-Hz contrast increased relative to surrogates that preserved the amplitude distribution and approximately matched the power spectrum. This pattern implicates temporal structure altered by the surrogate procedure and is compatible with both proposed mechanisms. Neighboring-block comparisons showed that the contrast persisted across adjacent film segments. The film also linked soundtrack category to visual and narrative content. Its speech annotations related word onsets to inferior frontal high-frequency power but showed no consistent relationship with STG sample entropy. Testing the mechanisms directly will require stimulus features measured on a common scale across speech and music.

Sample entropy connects these observations to studies of spontaneous activity at a slower timescale. Lower resting BOLD entropy has been associated with stronger task activation and deactivation (Lin et al., 2022). Hearing loss has also been linked to altered spontaneous STG activity, including lower resting BOLD sample entropy in a focal right STG region (Chen et al., 2018; Zhu et al., 2026). The intracranial results extend this question to faster local electrical fluctuations during listening. Separately acquired fMRI showed stronger bilateral STG activation during speech, while sample entropy estimated from short BOLD blocks showed little difference between conditions. Measuring both properties during rest and listening in the same individuals would help establish how spontaneous temporal organization and auditory history relate to responses to natural sounds.

During natural listening, speech elicited stronger high-frequency and BOLD responses in the human superior temporal gyrus. Yet it was during speech that similar patterns of local low-frequency activity were more likely to remain similar as they unfolded. Together, these findings give the speech preference of the STG a temporal dimension. Speech and music differ both in how strongly they recruit this shared auditory region and in how consistently its local activity follows recurring patterns over time.

## Methods

### Dataset and study design

We reanalysed the publicly released ds003688 film and iEEG-rest recordings (Berezutskaya et al., 2022a; Berezutskaya et al., 2022b). The source cohort comprises 51 participants with film iEEG, 30 with film fMRI and 18 with both modalities at different sessions; the analysis cohorts below are smaller because of region-specific electrode coverage and completeness requirements. In participants with both modalities, the source study acquired fMRI before iEEG, so they had already encountered the film at the time of intracranial recording (Berezutskaya et al., 2022a). The naturalistic film alternates approximately 30-s music and speech epochs. For the primary iEEG contrast, we retained the first six complete music and six speech windows from each participant’s film recording and excluded the seventh music window. The seven-music-versus-six-speech aggregate is retained as a sensitivity analysis. Rest iEEG was segmented into consecutive 30-s windows. The original source study describes consent, ethics approval, acquisition and anonymisation (Berezutskaya et al., 2022a).

The source study recorded intracranial EEG during film viewing and rest in patients undergoing clinical monitoring for epilepsy. Recordings were obtained from subdural or depth electrodes placed according to clinical needs. The BOLD data analysed here were acquired during viewing of the same film in separate 3 T MRI sessions, with a repetition time of 608 ms (Berezutskaya et al., 2022a).

### Electrode localisation and matching

We used the completed FreeSurfer reconstructions and source electrode/channel metadata to assign good ECoG or sEEG contacts to anatomical STG and IFG. Selection was restricted to the left hemisphere, the contact’s primary anatomical label, good status across the relevant recordings and availability in each compared condition. Within each participant and region, we selected the three eligible contacts closest to the reconstructed pial surface. Contacts and the number of 30-s blocks were then fixed across conditions. Population summaries first averaged windows and contacts within participant; no contact or block was treated as an independent participant. The six-band STG–IFG interaction required both regions and comprised 18 participants; the main STG-only analysis comprised 30 and IFG-only analysis 28.

### iEEG preprocessing and sample entropy

For the primary 1–10-Hz analysis, good intracranial channels were common-average referenced, filtered to 1–10 Hz and resampled to 25 Hz. Each native 30-s film block or matched rest window was analysed separately. Sample entropy was the negative natural logarithm of the conditional recurrence proportion for templates of length m + 1 relative to m, excluding self-matches (Richman and Moorman, 2000). We fixed m = 3 and r = 0.6 times the standard deviation of the respective analysed window. These parameters define the reported temporal-irregularity measure; absolute values should not be compared with SampEn from another sampling rate, representation or tolerance rule (Supplementary Table S12).

For the six-band follow-up, we computed SampEn of Hilbert amplitude envelopes in delta (1–4 Hz), theta (4–8 Hz), alpha (8–12 Hz), beta (13–30 Hz), low gamma (30–50 Hz) and HFB (60–100 Hz). The band-envelope series were resampled to 25 Hz and used the same m and window-relative r. Filtering and Hilbert transformation were applied with surrounding temporal padding before extracting the central 30 s. IAAFT surrogates were generated separately for each analysed window; the main 1–10-Hz analysis used five surrogates with 20 iterations, and the six-band follow-up used 20 surrogates. Excess SampEn was original minus mean surrogate SampEn. The surrogate construction approximately matches the observed window’s amplitude distribution and power spectrum (Schreiber and Schmitz, 1996) (Supplementary Tables S8, S9, S10, and S11).

### Spectral descriptors and stimulus annotations

Welch estimates supplied integrated band power and relative band power; a separate exploratory analysis estimated the height of local theta, alpha, beta and low-gamma spectral peaks above a robust fitted log-log aperiodic background. The peak metric is not a formal parameterisation of oscillatory bursts. HFB power was quantified separately and was not interpreted as a narrowband rhythm. The duration-aligned power/BEN join contained 17 participants because one participant exceeded a prespecified 0.05-s segment-duration mismatch; the spectral-peak analysis contained 18. We aligned publicly released word, phoneme, syllable, sentence and clause onset annotations to the same six 30-s speech blocks. Each annotation supplied an onset count per block; word intervals additionally supplied annotated-speech occupancy. For stimulus-feature inference, all 720 permutations of the six block labels were applied jointly across participants before averaging within-participant correlations. These annotations describe speech content and timing, not sound intensity or a music–speech acoustic continuum.

### fMRI processing and regional analyses

Thirty film BOLD runs were preprocessed with fMRIPrep 25.2.5. The primary activation analysis used a bilateral Harvard–Oxford STG maximum-probability mask thresholded at 25% and aligned to MNI152NLin2009cAsym 2-mm space. The first-level model included film-condition regressors convolved with a canonical haemodynamic response, 1/128-Hz high-pass filtering, AR(1) temporal noise, 24 motion expansions, up to six aCompCor components and expanded framewise-displacement outlier regressors. Group inference used participant-level music-minus-speech STG estimates and a model with mean framewise displacement; excluding three prespecified descriptive motion-flag cases formed a sensitivity analysis (Supplementary Table S15).

For BOLD SampEn, nuisance-regressed signals were analysed voxelwise before regional averaging. Each condition contributed six equal 49-sample blocks; blockwise estimates were averaged or template matches were pooled across blocks without constructing cross-block templates. Blockwise demeaning removes constant shifts, not time-varying responses. No resting-state fMRI run was available, and no fMRI-rest entropy comparison was made.

### Statistical analysis

The participant was the inferential unit throughout. Paired t and Wilcoxon tests summarised the main three within-region condition contrasts; Benjamini–Hochberg q values were computed separately for each region and metric across those three contrasts. Local symmetric film comparisons were averaged within participant before testing, and time-adjusted models used participant-clustered errors. The six-band STG– IFG interaction was [(STG music − speech) − (IFG music − speech)]; exact two-sided sign-flip probabilities and six-band Benjamini–Hochberg q values were calculated separately for original and excess SampEn. Spectral, word-annotation and cross-modal associations were exploratory. The common-stimulus annotation analysis permuted six film blocks jointly across participants rather than counting repeated observations of the same stimulus as independent.

## Data availability

The de-identified source data are available as OpenNeuro ds003688, version 1.0.7 (https://doi.org/10.18112/openneuro.ds003688.v1.0.7). The film itself is subject to the access conditions described by the dataset authors (Berezutskaya et al., 2022a).

## Notes

https://doi.org/10.18112/openneuro.ds003688.v1.0.7

### Competing Interest Statement

The authors have declared no competing interest.

### Summary of Updates

The previous version used the original title; the title has now been updated to match the text.

